# Rewiring of master transcription factor cistromes during high-grade serous ovarian cancer development

**DOI:** 10.1101/2023.04.11.536378

**Authors:** Robbin A. Nameki, Heidi Chang, Pak Yu, Forough Abbasi, Xianzhi Lin, Jessica Reddy, Marcela Haro, Marcos AS Fonseca, Matthew L. Freedman, Ronny Drapkin, Rosario I. Corona, Kate Lawrenson

## Abstract

The transcription factors MECOM, PAX8, SOX17 and WT1 are candidate master regulators of high-grade serous ‘ovarian’ cancer (HGSC), yet their cooperative role in the hypothesized tissue of origin, the fallopian tube secretory epithelium (FTSEC) is unknown. We generated 26 epigenome (CUT&TAG, CUT&RUN, ATAC-seq and HiC) data sets and 24 profiles of RNA-seq transcription factor knock-down followed by RNA sequencing in FTSEC and HGSC models to define binding sites and gene sets regulated by these factors in *cis* and *trans*. This revealed that MECOM, PAX8, SOX17 and WT1 are lineage-enriched, super-enhancer associated master regulators whose cooperative DNA-binding patterns and target genes are re-wired during tumor development. All four TFs were indispensable for HGSC clonogenicity and survival but only depletion of PAX8 and WT1 impaired FTSEC cell survival. These four TFs were pharmacologically inhibited by transcriptional inhibitors only in HGSCs but not in FTSECs. Collectively, our data highlights that tumor-specific epigenetic remodeling is tightly related to MECOM, PAX8, SOX17 and WT1 activity and these transcription factors are targetable in a tumor-specific manner through transcriptional inhibitors.

## INTRODUCTION

Ovarian cancer is one of the most lethal gynecologic malignancies with approximately 313,959 cases and 207,252 annual deaths globally (Sung et al., 2021). Epithelial tumors comprise ∼90% of ovarian cancer cases. The most common subtype, high-grade serous ‘ovarian’ cancers (HGSCs) are characterized by a 5-year survival rate of 30% for those diagnosed with advanced-stage disease - which is a majority of cases (NIH, n.d.). Recent therapeutic advances including angiogenesis and poly ADP-ribose polymerase (PARP) inhibition have improved progression-free survival rates for this disease (Garcia et al., 2020; Moore et al., 2018). However, an improved understanding of the non-genetic (epigenetic) landscapes of HGSC remains a research priority for the development of additional approaches for personalized therapeutics - “precision medicine” - for ovarian cancer patients (Kuroki and Guntupalli, 2020).

In other tumor types, epigenome profiling studies have coupled the activity of critical transcription factors regulators to transcriptional fingerprints of tumors during initiation, progression, drug resistance and trans-differentiation (Baca et al., 2021; Bell et al., 2019; Chi et al., 2019). ‘Master’ transcription factors are lineage-enriched developmental regulators that are often functionally co-opted during tumorigenesis to support atypical cell survival, metastasis or drug resistance and therefore represent an important source of therapeutic targets for cancer (Garraway and Sellers, 2006). Until recently, the critical TFs driving HGSC development were poorly characterized. In the last few years, our laboratory and others have identified and validated MTFs in HGSC, including PAX8, SOX17 and MECOM (Bleu et al., 2021; Lin et al., 2022; Reddy et al., 2021). PAX8 and SOX17 in particular cooperate to positively regulate cell cycle progression and angiogenesis in HGSC (Chaves-Moreira et al., 2022; Lin et al., 2022; Reddy et al., 2021; Shi et al., 2019). However, the activity of these TFs in benign precursor cells - fallopian tube secretory epithelial cells (FTSECs), is not fully understood. Here, we aimed to investigate our hypothesis that common factors dominate control of lineage-specific transcriptional programs in precursor fallopian tube secretory epithelium and that these regulatory networks are reprogrammed to promote pro-oncogenic signaling in HGSC.

## RESULTS

### Lineage-enrichment of MECOM, PAX8, SOX17 and WT1 in pan-normal and cancer tissues

High level co-expression of MECOM, PAX8, SOX17 and WT1 characterizes HGSCs compared to other tumor types (Reddy et al., 2021). To ask whether this lineage-restricted expression pattern is recapitulated in benign tissues, we systematically compared expression patterns of these factors in the Genotype-Tissue Expression (GTEx) and the Cancer Genome Atlas (TCGA) databases. K-means clustering based on *MECOM, PAX8, SOX17* and *WT1* mRNA expression in 21 normal tissues revealed 5 main clusters (**Figure 1A**). A distinct cluster consisting of fallopian tube and uterus tissues was characterized by *MECOM, PAX8, SOX17* and *WT1* expression (MTF^high^ cluster). A majority 57% (12/21; MTF^low^ cluster) of normal tissue types did not highly express any of these four transcription factors and 33% (7/21; M^high^, M&P^high^ cluster and SOX17^high^ cluster) expressed just one or two candidate MTFs. *WT1* was expressed in both benign fallopian tube and uteri, but in tumors, was only observed in HGSC and not uterine adenocarcinoma (**Figure 1A**). Positively correlated expression between these four factors is indicative of a core regulatory circuitry characterized by high-level expression of each factor. Fallopian tube tissues showed the greatest correlations of *MECOM, PAX8, SOX17* and *WT1* expression compared to all other normal tissue types (Ranked 1/21, Pearson correlations, **Figure 1B**, **Supplementary Table 1**). The top 3 tumor tissues from TCGA were represented by gynecological malignancies including CESC (1/27), UCS (2/27) and HGSC (3/27) (**Figure 1B**, **Supplementary Table 2**).

**Figure 1.**
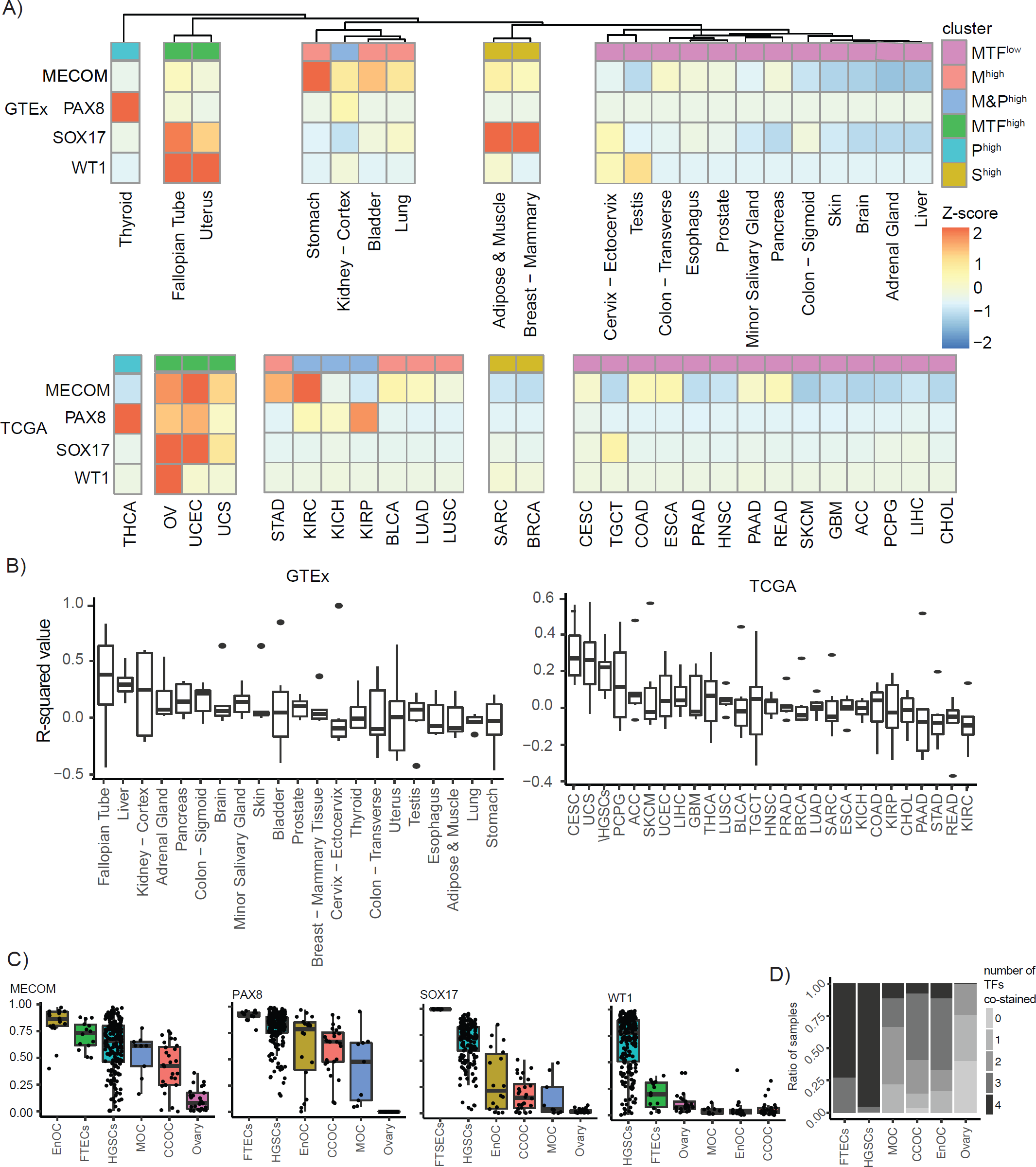
mRNA and protein expression of MECOM, PAX8, SOX17 and WT1 loci in normal tissues and cancer. (A) Unsupervised hierarchical clustering of *MECOM, PAX8, SOX17* and *WT1* mRNA expression in the pan-normal GTEx dataset. TCGA data was clustered based on the 5 main clusters of TF expression from GTEx. MTF^low^ = Cluster of tissues that did not or lowly expressed MTFs, MTF^high^ = cluster of tissues that moderately or highly expressed all four MTFs, M^high^ = Cluster of tissues that highly expressed MECOM, M&P^high^ = cluster of tissues that highly expressed both MECOM and PAX8, P^high^ = cluster of tissues that expresses PAX8 and S^high^ = cluster of tissues that expresses SOX17). (B) A boxplot of the average Pearson correlation values of MECOM, PAX8, SOX17 and WT1. Ranked from highest average Pearson correlation value to lowest. (C) A boxplot representing the mean positivity rate of MECOM, PAX8, SOX17 and WT1 expression in each histotype. In B and C, the limits of the boxes represent the interquartile range, and the limits of error bars represent the minimum and maximum value without outliers (D) Ratio of samples with number of co-stained TFs based on a 0.1 positivity rate threshold.

Epithelial ovarian cancers comprise multiple histologic subtypes, with diverse cells-of-origin (Cochrane et al., 2017). MECOM, PAX8, SOX17 and WT1 protein expression was quantified in high-grade serous (n = 181-200), clear cell (n = 27-28), endometrioid (n = 18) and mucinous (n = 9) ovarian cancers *via* immunohistochemistry. Benign ovarian cortex (n = 25) and fallopian tube (n = 12-14) tissues were also evaluated as controls. As previously reported, PAX8 (0.44-0.82 mean positivity rate) and SOX17 (0.10-0.99 mean positivity rate) expression was pervasive in serous, endometrioid and clear cell tumors, with heterogenous expression in mucinous tumors (**Figure 1C**, **Supplementary Table 3**) (Dinh et al., 2021; Lin et al., 2022). MECOM (0.44-0.82 mean positivity rate) exhibited a similar expression pattern to PAX8 and SOX17 **(Supplementary Table 3)**. 96% of HGSCs (0.68 mean positivity rate; standard deviation, 0.27), and 86% of fallopian tube tissues (0.19 mean positivity rate; standard deviation, 0.13) expressed WT1, but this marker was less commonly expressed in non-high-grade serous subtypes (Acs et al., 2004; Köbel et al., 2008; Shimizu et al., 2000) **(Table 1)**. A majority of FTSECs (8/11) and HGSCs (142/149) co-stained for all 4 transcription factors (**Figure 1D**, **Supplementary Table 3**). Together, these data indicate that co-expression of *MECOM, PAX8, SOX17* and WT1 is a unique characteristic of HGSC and fallopian tube precursor tissues.

### Super-enhancer landscape of MECOM, PAX8, SOX17 and WT1 in FTSECs and HGSCs

Super-enhancer association is a prominent feature of MTFs that contributes to the maintained high expression of MTFs in a lineage-enriched manner (Bradner et al., 2017; Chapuy et al., 2013; Chen et al., 2020; Eliades et al., 2018; Jiang et al., 2020; Sanda et al., 2012). We interrogated global landscapes of the active chromatin marked by H3K27ac in FTSECs and HGSCs, including ChIP-seq and newly generated CUT&TAG profiles of primary tissues, cell lines and *ex vivo* cultures (Corona et al., 2020). *MECOM, PAX8, SOX17* and *WT1* were associated with super-enhancers in FTSECs and HGSCs (**Figure 2A**). Inspection of constituent enhancers revealed a marked remodeling of the SOX17 super-enhancer in HGSCs and an acquired acetylation of one of the MECOM promoters (**Figure 2A**). High-throughput chromosome conformation capture (Hi-C) was conducted in 2 FTSEC (FT246 and FT282) and 2 HGSC (KURAMOCHI and OVCAR4) cell lines to measure the contact loops associated with enhancers in these regions. Hi-C loops were often anchored to constituent enhancers that were part of the super-enhancer regions in FTSECs and HGSCs (**Figure 2C**). These data suggest that the *MECOM, PAX8, SOX17* and *WT1* are super-enhancer associated TFs in FTSECs and HGSCs that are likely contributing to the maintained lineage-enriched expression of these TFs from FTSEC to HGSC development.

**Figure 2.**
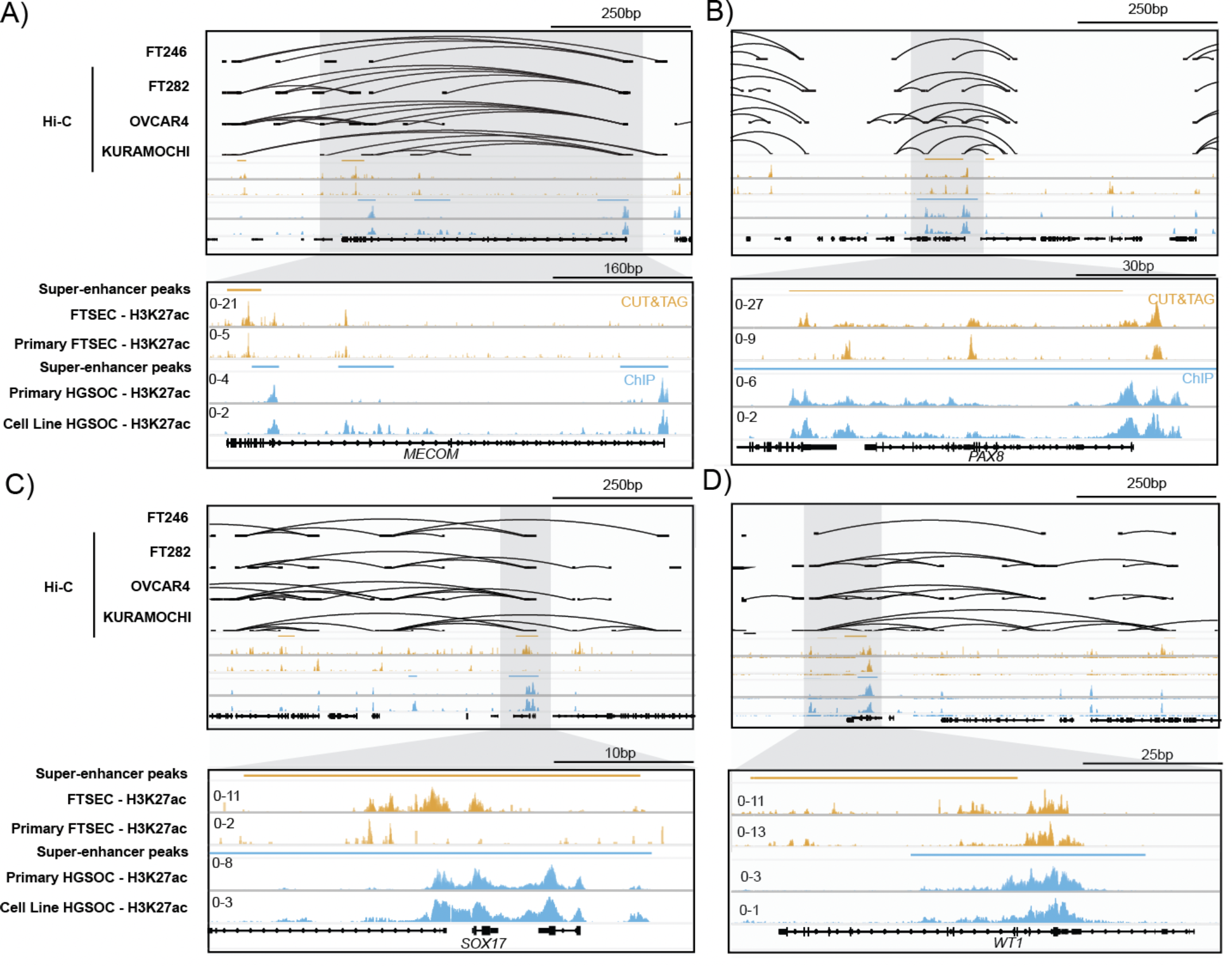
MECOM, PAX8, SOX17 and WT1 super-enhancer landscape in FTSECs and HGSCs. Landscapes of active chromatin and chromatin loops (based on Hi-C maps) in FTSECs and HGSCs. (A) *MECOM,* (B) *PAX8,* (C) *SOX17* and (D) *WT1*.

### Tumor-specific effects of genetic and pharmacologic master TF depletion

Cancer cells are often addicted to the expression of MTFs and MTF depletion leads to tumor cell death (Chapuy et al., 2013; Jiang et al., 2020; Witwicki et al., 2018). TF expression was potently depleted in 2 FTSEC (FT246 and FT282) and 2 HGSC (KURAMOCHI and OVCAR4) cell lines using siRNA (**Supplementary Table 4**). MECOM, PAX8 and SOX17 were indispensable for HGSC survival as previously reported (**Figure 3A & 3B**) (Reddy et al., 2021). WT1 was also necessary for HGSC survival. By contrast, FTSECs were dependent on PAX8 and WT1 but not MECOM and SOX17 (**Figure 3A & 3B**).

**Figure 3.**
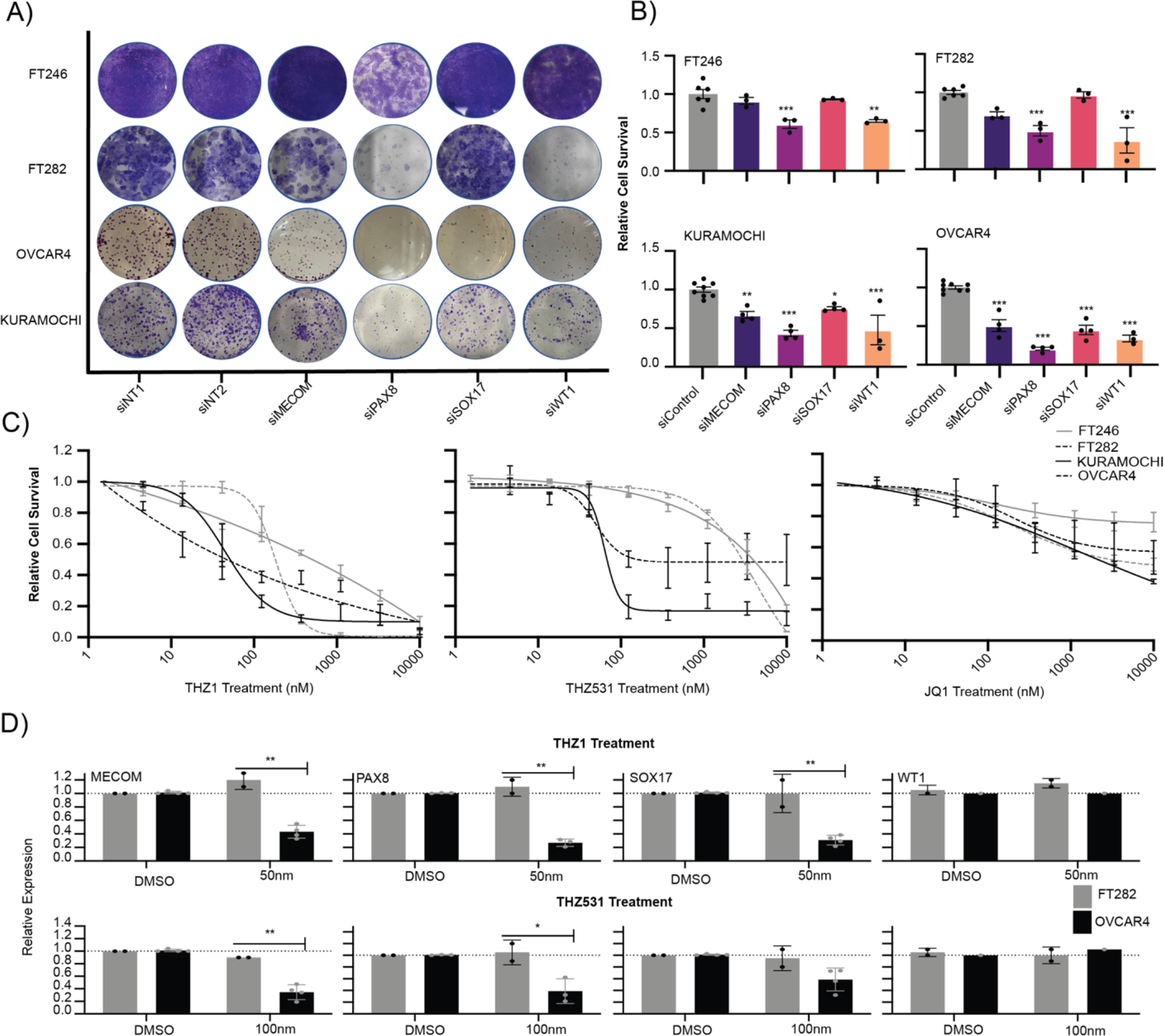
MECOM, PAX8, SOX17 and WT1 are tumor-specific therapeutic targets. (A) TF knock-down followed by colony formation assays stained with crystal violet. Representative wells are shown. (B) Barplots representing the quantification of crystal violet staining in Figure 3A. Error bars represent biological replicates. (C) Dose response curves for FT246, FT282, KURAMOCHI and OVCAR4 cells treated with THZ1, THZ531 and JQ1. Error bars represent standard deviation of mean cell survival values from biological replicates. (D) RT-qPCR quantification of *MECOM, PAX8, SOX17* and *WT1* expression upon THZ1 and THZ531 treatment in FT282 and OVCAR4 cells. Data for *MECOM, PAX8* and *SOX17* expression upon THZ1 and THZ531 treatment of OVCAR4 cells are reproduced from Reddy et al., 2021.

Cyclin-dependent kinase 7/12/13 inhibitors are hypothesized to have anticancer effects *via* disruption of the transcriptional initiation machinery that in enriched in super-enhancers regulating MTFs and other lineage-specific genes (Chapuy et al., 2013; Chen et al., 2020; Jiang et al., 2020). OVCAR4 cells were sensitive to THZ1 (CDK7 inhibitor) and THZ531 (CDK7/12 inhibitor) treatment as previously reported (THZ1 IC_50_ = 44.93nM; n=4, THZ531 IC_50_ = 63.42nM; n=4), a trend which we replicated in KURAMOCHI cells (THZ1 IC_50_ = 14nM; n=5, THZ531 IC_50_ = 48.90nM; n = 2)(**Figure 3C**) (Reddy et al., 2021). FT246 (THZ1 IC_50_ = 319.70nM; n = 3, THZ531 IC_50_ =5033nM, n = 3) and FT282 (THZ1 IC_50_ = 183.9nM; n = 3, THZ531 IC_50_ = 4135nM; n =3) cells were 8 and 82 times more resistant for THZ1 and THZ531 treatment respectively, suggesting a tumor-specific mechanism for the anti-proliferative effect of these drugs (**Figure 3C**). Expression of *MECOM, PAX8* and *SOX17* was downregulated by 30-75% (P < 0.007, multiple comparison test) in HGSCs treated with short-term (6 hour) treatment of THZ1 at IC_50_. By contrast, in FTSECs, expression of all four TFs was unaffected by exposure to IC_50_ THZ1 treatment (**Figure 3D**). Context-specific inhibition of MTFs was also observed for THZ531, in which the expression of *MECOM* and *PAX8* was 60% lower in OVCAR4 compared to expression in FT282 upon THZ531 treatment at IC_50_. WT1 was not sensitive to THZ1 or THZ531 treatment in either FTSECs nor HGSCs. These data suggest that the antiproliferative effects of THZ1 and THZ531 in HGSC cells may be due to tumor-specific inhibition of *MECOM, PAX8* and *SOX17* expression by these drugs.

### MECOM, PAX8, SOX17 and WT1 forms a complex core-regulatory circuit that is rewired during HGSC development

We dissected the DNA binding pattern of MECOM, PAX8, SOX17 and WT1 in FT282 and KURAMOCHI cells using Cleavage Under Targets and Release using Nuclease (CUT&RUN) (Skene et al., 2018). All four TFs co-occupied the regulatory elements within each TF’s loci in both cellular contexts, further corroborating the involvement of these TFs in autoregulation plus expression of the other MTFs (**Figure 4A**). MTFs are expected to co-occupy active enhancer regions to control global gene expression programs, consistent with this MECOM, SOX17 and WT1 binding was enriched within PAX8 bound regions in both cellular contexts (**Figure 4B & Figure 4C**). These regions were associated with active enhancer regions marked by H3K27ac, but did not coincide with heterochromatin regions marked by H3K27me3 (**Figure 4B & Figure 4C**). In a set intersection analysis of all available TF peaks, the proportion of TF peaks co-bound by at least two master transcription factors was 11% higher in HGSC compared to FTSEC (54%, 32,122 out of 59,349 for FT282; 65%, 37,724 out of 58,000 peaks for KURAMOCHI) (**Figure 4D**). Furthermore, the number of regions co-bound by all 4 TFs was around 50% higher in KURAMOCHI compared to FT282 (14,547 and 10,588 peaks respectively). Based on genomic and histone context, we annotated the TF peaks as binding within promoters, typical enhancers, super-enhancers, typical methylated regions, and super-methylated regions. Regions bound by all 4 TFs were most frequently defined as typical enhancers (63.7% for FT282; 45.7% for KURAMOCHI), or super-enhancers (10.9% for FT282; 8.95% for KURAMOCHI) and were rarely methylated (typical methylated regions - 0.9% for FT282; 0.72% for KURAMOCHI; super-methylated regions - 0.22% for FT282; 0.082% for KURAMOCHI)(**Figure 4E**). These data indicate that MECOM, PAX8, SOX17 and WT1 preferentially co-occupy active enhancers across the genome of FTSECs and HGSCs alike.

**Figure 4.**
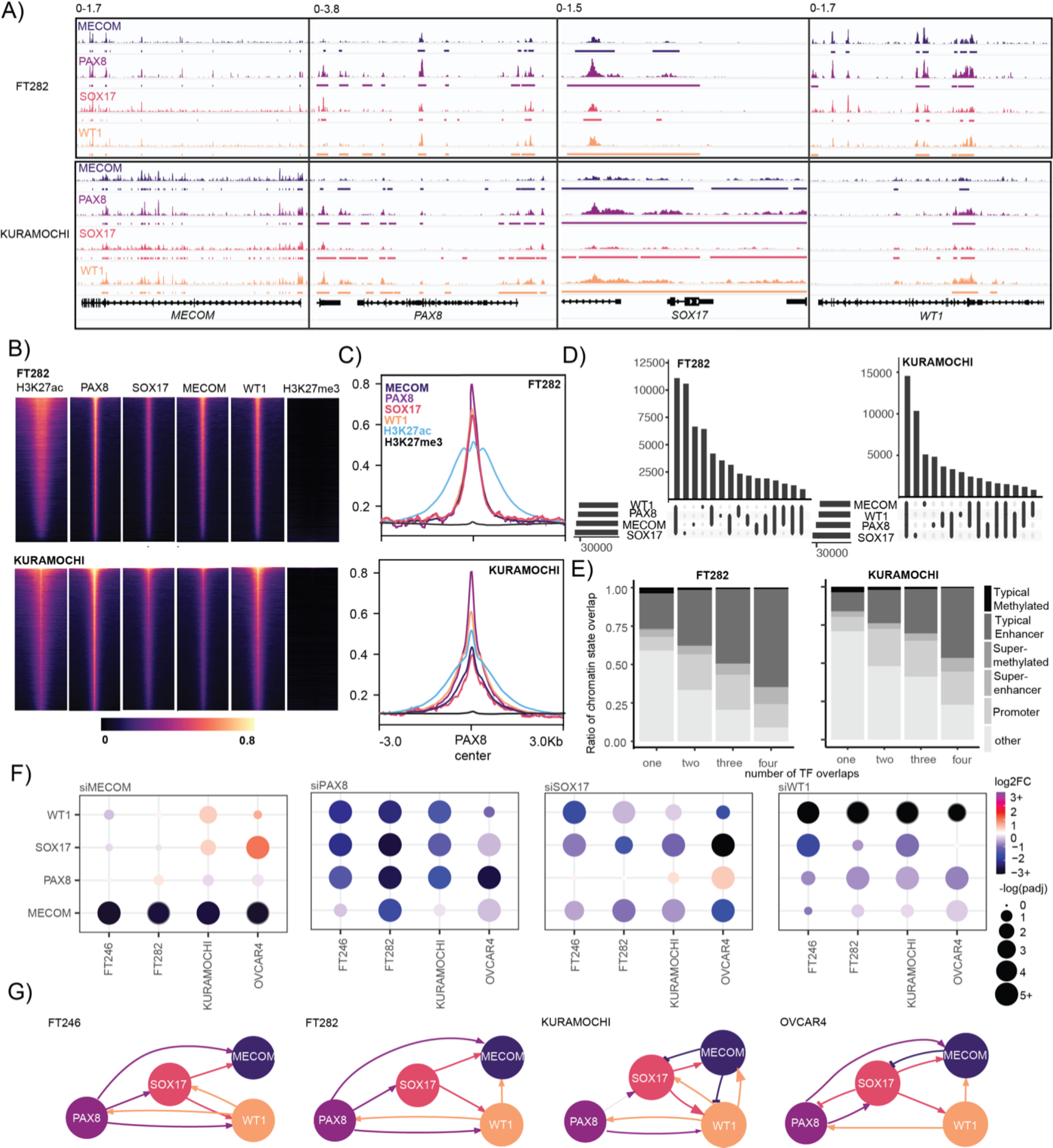
MECOM, PAX8, SOX17 and WT1 form a core-regulatory circuit in FTSEC to HGSC but cross-relation between factors changes during tumorigenesis. (A) MECOM, PAX8, SOX17 and WT1 co-occupies its own and others genomic loci. (B) MECOM, PAX8, SOX17 and WT1 co-occupy active enhancer regions across the genome. CPM-normalized CUT&RUN reads were centered on 3 kilobase windows of FTSEC or HGSC-specific PAX8 peaks. Rows are the same across feature. (C) Metagene plot, MECOM, PAX8, SOX17, WT1, H3K27ac and H3K27me3 signal centered on FT of HGSC-specific PAX8 peaks. (D) Set analysis of CUT&RUN peaks from representative MECOM, PAX8, SOX17 and WT1 samples in FT282 and KURAMOCHI. (E) Chromatin state of TF peaks categorized by number of TF overlaps. (F) MECOM, PAX8, SOX17 and WT1 co-regulation based on TF knock-down followed by RNA-seq and differential expression analysis with DESEQ2. (G) Node and edge plot representing co-regulation of each TF based on gene expression measured by RNA-seq after TF knock-down.

At the transcriptional level, PAX8 knock-down led to the depletion of *MECOM, SOX17* and *WT1* expression in both FTSEC and HGSC lines (**Figure 4F**, **Supplementary Table 4**). WT1 knock-down also led to the depletion of *MECOM, PAX8* and *SOX17*. Depletion of SOX17 resulted in downregulation of *WT1* and *MECOM* expression, and a reproducible trend towards up-regulation of PAX8 in HGSC models (**Figure 4F**, **Supplementary Table 4**). MECOM depletion resulted in upregulated SOX17 expression in HGSC cells as previously reported (Reddy et al., 2021) and also upregulated WT1 expression in HGSC cells (**Figure 4F**, **Supplementary Table 4**). In summary, patterns of cross-regulation between these transcription factors is altered in HGSCs compared to FTSECs, with PAX8 playing a dominant role in both contexts but MECOM and SOX17 participating more in the circuitry in tumor cells compared to precursor cells (**Figure 4G**).

### The master transcription factor cistrome is rewired between FTSEC to HGSC development

Comparison of TF CUT&RUN binding sites between FT282 and KURAMOCHI showed that for each factor, only 15-23% of peaks overlap between the two cellular contexts (**Figure 5A**). For example - 18,488 PAX8 peaks were specific to FTSECs, 20,717 were specific to HGSC and 14,544 were in common to the two contexts. In FTSECs, FTSEC-specific and common PAX8 peaks exhibited higher local chromatin activity (H3K27ac positivity) and were more accessible than the HGSC-specific peaks. Remarkably, SOX17, MECOM and WT1 co-localized to the FTSEC-specific and common PAX8 bound regions and were depleted at HGSC-specific peaks (**Figure 5B**). The inverse pattern was observed for HGSC-specific PAX8 peaks, which were active and more accessible in HGSC cells compared to in FTSECs, and co-occupied by WT1, SOX17 and MECOM in HGSC specifically. TF binding sites specific to FT282 or KURAMOCHI were enriched in typical enhancers or super-enhancers, but not promoters (**Figure 5C**).

**Figure 5.**
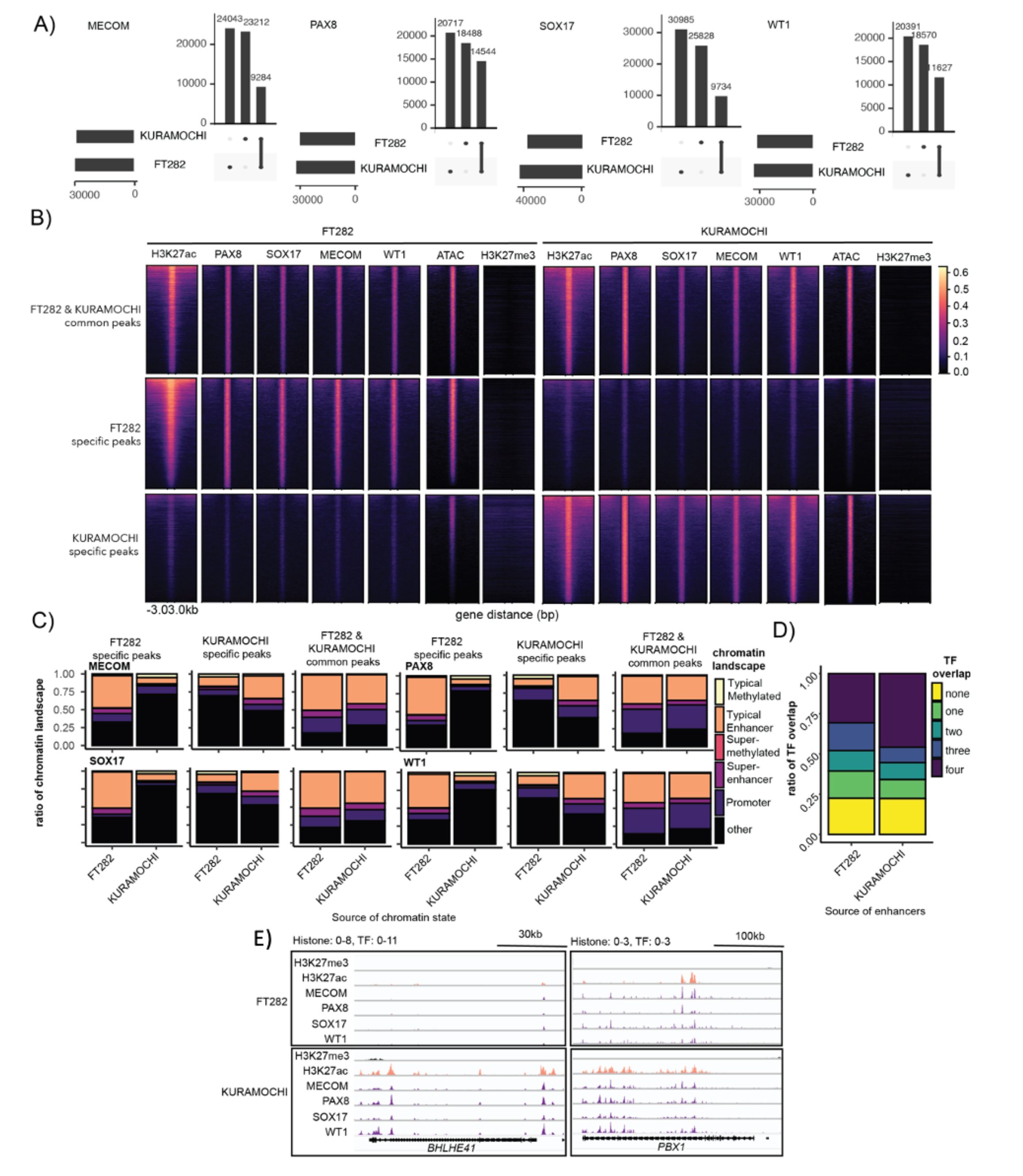
MECOM, PAX8, SOX17 and WT1 cistromes are remodeled during tumorigenesis. (A) Set analysis of TF binding sites that were common or specific to FT282 or KURAMOCHI. (B) MECOM, PAX8, SOX17 and WT1 co-occupies regions that were bound in a common or context-specific manner. CPM-normalized CUT&RUN reads were centered on 3 kilobase windows of PAX8 peak start and stop positions. Rows are the same across samples. (C) Ratio of chromatin states associated with TF binding regions categorized by cellular context. (D) Ratio of FT282 and KURAMOCHI specific enhancers bound by one, two, three or four TFs. (E) *BHLHE41* and *PBX1* loci displaying the co-localization of MECOM, PAX8, SOX17 and WT1 at a KURAMOCHI specific super-enhancer.

We sought to contextualize these multi-bound TF binding sites against the background of active enhancers gained or lost during HGSC development. Enhancers specific to FT282 or KURAMOCHI were more likely to be bound by TFs compared to the null distribution (P < 1.0×10^−4^). A large proportion (3068/6703 FT282 specific enhancers and 3017/9879 KURAMOCHI specific enhancers) were co-bound by all four TFs, confirming that context specific enhancers are characterized by the binding of MECOM, PAX8, SOX17 and WT1 (**Figure 5D**). Analysis to identify HGSC specific super-enhancer associated genes bound by MECOM, PAX8, SOX17 and WT1 revealed two other candidate master regulators of HGSCs, *BHLEH41* and *PBX1* (**Figure 5F**). Overall, although the core-regulatory circuitry of MECOM, PAX8, SOX17 and WT1 are maintained between FTSECs and HGSCs, HGSCs acquire tumor-specific TF DNA binding sites through extensive remodeling of cooperative binding events involving these four factors.

### Master transcription factors acquire target genes during HGSC development

RNA-seq after TF knockdown were integrated with TF CUT&RUN to contrast the cistromes of MECOM, PAX8, SOX17 and WT1 in FTSECs and HGSCs. ERCC spike-in normalization was performed to enable sensitive detection of global transcriptional changes (Lovén et al., 2012). PAX8 preferentially functioned as a transcriptional repressor in tumor and normal cells alike. The dominant effect of MECOM depletion was gene down-regulation, suggesting this factor mainly functions as a transcriptional activator, whereas depletion of SOX17 was associated with a relatively equal number of up- and down-regulated differentially expressed genes (DEGs), implying this gene both activates and represses gene expression. WT1 displayed the most marked context-specific activity as in FTSECs this factor functioned almost exclusively as a transcriptional activator and dominated control of the transcriptome whereas in HGSC cells WT1 regulated ∼50% fewer genes and switched to largely function as a transcriptional repressor (**Figure 6A & B, Supplementary Figure 1**).

**Figure 6.**
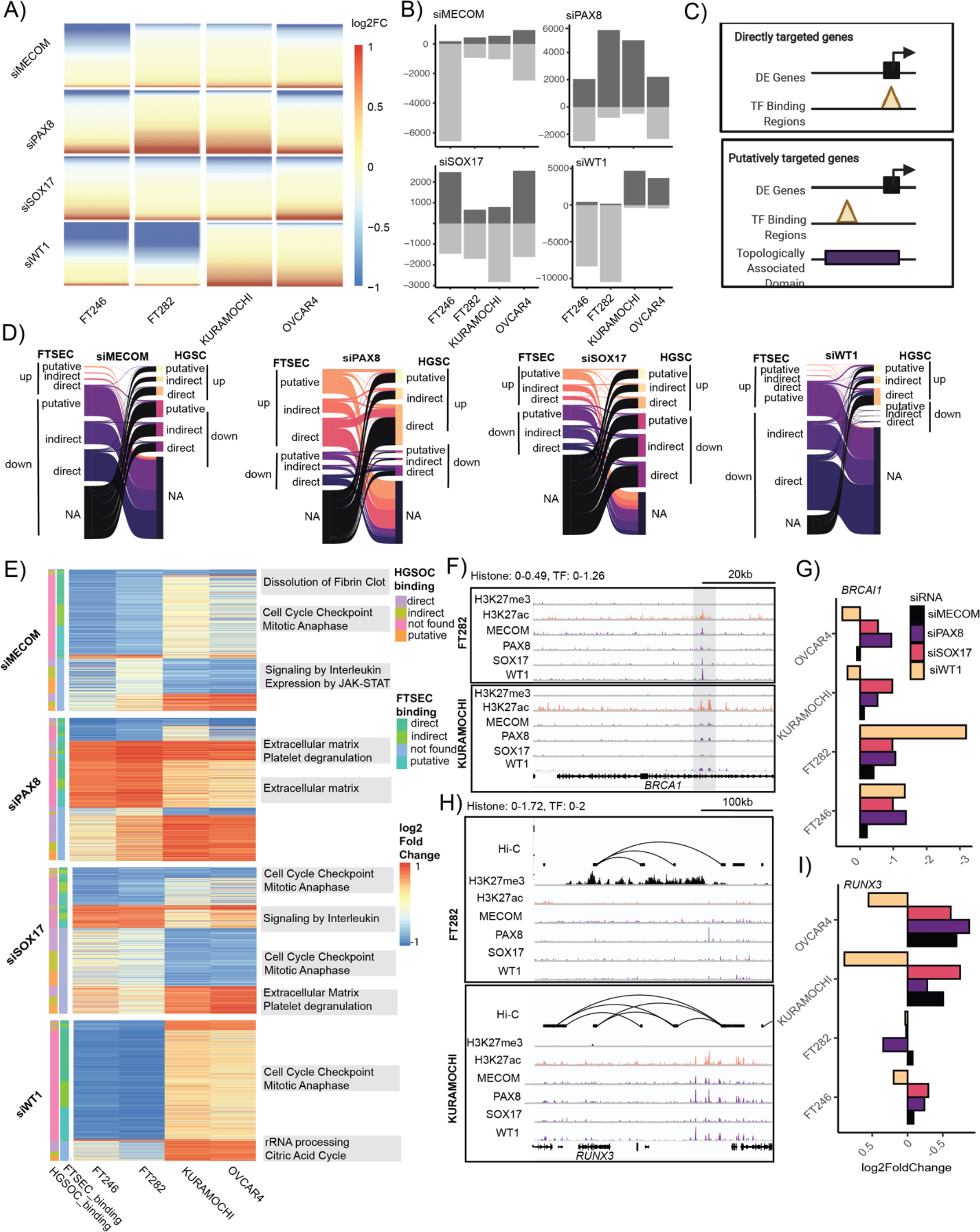
Gene regulation by MECOM, PAX8, SOX17 and WT1 in FTSECs and HGSC. (A) Log_2_ fold-change of 28,158 genes following normalization informed by ERCC spike-in RNA content, (B) Number of differentially expressed genes for each TF knock-down based on absolute log_2_ fold-change ≥ 0.5. (C) Schematic to integrate differential expression, TF binding and topologically associated domain (TAD) maps. (D) Alluvial plot displaying the status of high confidence differentially expressed genes following TF depletion in FTSECs and HGSCs. (E) Heatmap and pathway analysis of genes displayed in D. (F) *BRCA1* locus highlighting H3K27ac signal and TF binding at the *BRCA1* promoter. (G) Log_2_ fold change of *BRCA1* expression following MECOM, PAX8, SOX17 and WT1 knock-down relative to scrambled controls. (H) Chromatin landscape of *RUNX3* locus in FTSECs and HGSCs. (I) Log_2_ fold change of *RUNX3* expression following MECOM, PAX8, SOX17 and WT1 knock-down relative to scrambled controls (RNA-seq).

High-confidence target genes for each factor (absolute log_2_FC ≥ 0.5 in both cell lines with TF-targeting siRNA versus two controls) were integrated with TF CUT&RUN to link TF binding events to differentially expressed genes within the same topologically associated domains (informed by Hi-C performed in the same cell lines) (**Supplementary Tables 5 & 6**, **Figure 6C**). Using this approach, we identified (1) direct TF target genes defined as DEGs with TF binding on the gene promoter, (2) putative TF target genes defined as TF binding within non-promoter regions within the same TAD as DEGs following TF knockdown and (3) indirect TF targets defined as DEGs with no TF binding within the same TAD. This identified 442, 1,397, 339 and 5,411 high-confidence direct or putative target genes for MECOM, PAX8, SOX17 and WT1 respectively in FTSECs and 279, 1,310, 578 and 1,494 direct or putative TF target genes in HGSCs (**Supplementary Table 7**). The majority of high confidence differentially expressed genes were FTSEC or HGSC specific across all TFs that were either directly, putatively or indirectly regulated (**Figure 6D**). Supervised clustering of differentially regulated genes revealed that 1,119 target genes repressed by WT1 in HGSCs were enriched in ribosomal RNA processing and citric acid cycle pathways. Of these, 612, 256 and 252 are either directly, putatively or indirectly regulated by WT1 respectively (**Figure 6E**). In addition, SOX17 acquired 501 down-regulated and 237 up-regulated directly, putatively or indirectly regulated genes that converged on cell cycle and extracellular matrix/platelet degradation pathways, respectively. *BRCA1,* the human tumor suppressor gene that is germline or somatically mutated in 20-25% of HGSCs (Manchana et al., 2019), is directly regulated by all four TFs in FTSECs. PAX8 and SOX17 maintain positive regulation of *BRCA1* in HGSCs, but regulation by MECOM and WT1 is lost in tumors, accompanied by a concomitant loss of WT1 and MECOM binding in an intronic regulatory element (**Figure 6 F,G**). A tumor specific enhancer co-occupied by TFs makes a contact loop spanning a distance of 300kb to the HGSC oncogene *RUNX3* (Lee et al., 2011; Nevadunsky et al., 2009)(**Figure 6H**). Furthermore, *RUNX3* is marked by the repressive marker H3K27me3 with little TF activity in FTSECs, suggesting this region undergoes a demethylation process during tumorigenesis (**Figure 6H**). TF knock-down followed by RNA sequencing data confirmed that MECOM, PAX8 and SOX17 putatively regulates *RUNX3* in HGSC lines (**Figure 6I**). Thus, the target genes of TFs are reprogrammed between FTSECs and HGSCs, resulting in the dysregulation of key pathways and genes involved in HGSC development.

## DISCUSSION

Often, maintained expression of lineage-restricted TFs is critical for tumor cell survival, making this class of transcription factors an “Achilles heel” for many tumor types. For example, the melanocyte identity and specification transcription factor MITF is an amplified oncogene that exhibits cell-type specific cellular dependencies in cutaneous melanoma (Eliades et al., 2018; Levy et al., 2006) and TFs involved in T-cell development such as TAL1, GATA3, RUNX1 and MYB form a core-regulatory circuit and are essential for the survival of T cell acute lymphoblastic leukemias (Mansour et al., 2014; Sanda et al., 2012). The functional role of these transcription factors become redirected during tumor development to support neoplastic transformation (Leach et al., 2017; Pomerantz et al., 2015). We and others have previously identified MECOM, PAX8 and SOX17 as lineage-defining master transcription factors in high-grade serous ovarian cancer (Bleu et al., 2021; Chaves-Moreira et al., 2022; Lin et al., 2022; Reddy et al., 2021), a particularly aggressive histological subtype of ovarian cancer associated with five-year survival rates under 50 per cent. While pervasively expressed in benign FTs and HGSCs, phenotypic roles, binding sites and target genes for these factors are altered in tumor cells compared to precursor cells. MECOM, PAX8, SOX17 and WT1 co-bind to over 10,000 HGSC-specific DNA binding sites, with a notable example being the co-localization of all four transcription factors at the *BHLHE41* and *PBX1* loci, HGSC-specific active regions bound by TFs that were not active in FTSECs. Critically, context-specific enhancers were nearly all bound by PAX8, SOX17, MECOM and WT1, suggesting that these four factors dominate control of epigenome reprogramming in HGSC. We were not able to elucidate in this study whether the genomic regions that acquire HGSC-specific TF binding sites are inherently at a primed or neutral state in FTSECs. Future studies using marks such as H3K4me1 will further delineate the epigenetic remodeling and alteration in chromatin states associated with these TF binding regions.

The transcriptome regulated by MECOM, PAX8, SOX17 and WT1 were also extensively reprogrammed between FTSECs and HGSCs. SOX17 gains 738 genes involved in cell cycle, extracellular matrix and platelet degranulation pathways. Platelet degranulation is an essential step for angiogenesis and this is consistent with previous findings that SOX17 cooperatively orchestrates angiogenic programs with PAX8 (Chaves-Moreira et al., 2022). WT1 function in HGSC is poorly understood, this study found that WT1 dominates control of the transcriptome in FTSECs but not in HGSC where it acquires the inhibition of genes involved in ribosomal RNA processing and citric acid cycle pathways. This extensive WT1 transcriptomic reprogramming may be critical for the switch from a FTSEC to a HGSC cell state.

While providing key insights into epigenome restructuring during HGSC development, this analysis raises many questions. For example, it is not yet known if one of the four factors possesses pioneer activity and nucleates the new enhancers acquired during tumorigenesis. Since related PAX family members can bind to and open heterochromatin (Pelletier et al., 2021), it is plausible that PAX8 may play a similar role. In addition, it is not clear how the same proteins acquire thousands of tumor-specific genomic binding sites while losing a set of normal tissue-specific binding sites, and whether these changes occur in a stepwise, gradual manner, or rapidly. None of these factors are frequently mutated in HGSCs, although PAX8 and MECOM are commonly amplified which may serve to reinforce and stabilize sustained high-level transcription at these loci. Substantial differences in absolute expression of WT1 protein between normal fallopian tube and HGSC that could potentially impact the stoichiometry of transcriptional complexes resulting in transcriptional circuit rewiring. Post-transcriptional or post-translational mechanisms may also play a role in modifying the activities of these four factors or critical cofactors.

Annotation of TF transcriptomes integrated with TF CUT&RUN revealed that WT1 positively regulates *BRCA1* in normal FTSECs but not in tumor cells, suggesting loss of WT1 binding at this locus may represent an alternative path to loss of *BRCA1* expression. Considering how patients with loss of *BRCA1* function are associated with increased sensitivity to chemotherapy and better overall survival (Bolton et al., 2012), continued efforts to characterize the roles of these factors in epigenetic regulation of homology-directed repair genes may reveal subsets of patients that may also benefit from PARP inhibitor therapy.

While we drew upon epigenome data sets generated in primary tissues and performed many assays in multiple *in vitro* models, the main limitation in this study lies in the reliance on a single prototype FTSEC and HGSC model for the TF binding profiling. Future studies performed in larger sample sets will be required to refine the TF binding patterns presented here. Nonetheless, the context specific molecular profiles and functional roles of these four factors opens up myriad new opportunities for ‘precision medicine’ for HGSC, perhaps most rapidly through the use of general transcriptional inhibitors, including those currently in Phase I clinical trials for solid tumors (Syros Pharmaceuticals, 2022).

## METHODS

### TCGA and GTEx expression analysis

Pan-cancer TCGA RNA sequence level 3 data was downloaded and curated as described previously (Reddy et al., 2021). Pan-normal GTEx RNA sequence data were downloaded and annotated from the GTEx portal (GTEx Analysis V8; https://www.gtexportal.org/home/datasets). Pearson correlation analysis was conducted by implementing the corr.test() function available from the Psych package in R.

### Immunohistochemistry & image analysis

MECOM immunohistochemistry was conducted by the Cedars-Sinai Cancer Biobank and Translational Research Core. Embedded slides were paraffinized at 72°C with EZ solution (Roche, Ventana) and MECOM antigen was retrieved with CC2 pre-diluted Tris solution for 64 minutes at 91℃. MECOM (Cell Signaling, 1:500, AB_2184098) primary antibodies were diluted in Antibody Dilution Buffer (AF1924, R&D), and treated for 32 minutes at 37℃, respectively. MECOM antibodies were detected with an Envision anti-Rabbit polymer (DAKO) for 32 minutes at room temperature. PAX8 and WT1 immunohistochemistry was conducted by the department of Pathology and Laboratory Medicine Division of Anatomic Pathology at Cedars-Sinai Medical Center. Pathology-grade, pre-diluted PAX8 (Cell Marque, no RRID available) and WT1 (Roche, no RRID available) antibodies were detected with the Ultraview DAB detection kit on the Ventana Benchmark Ultra.

Digital image analysis (Qupath) was used to determine the percent of epithelial cells in each sample that stained positive for each transcription factor (percent positivity rate), and the proportion of each tumor type that co-expresses all four factors was determined by measuring for samples with a mean positivity of 0.1 or higher.

### Cell culture

Only human cell lines were used in this study and were regularly authenticated using STR profiling, performed at the University of Arizona Genetics Core. FT246 and FT282 cells were cultured in DMEM/F12 with 10% fetal bovine serum and 1× penicillin/streptomycin. OVCAR4 cells were cultured in RPMI 1640 supplemented with 10% bovine serum, 1× nonessential amino acid (NEAA), insulin (11.4μg/ml) and 1× penicillin/streptomycin. KURAMOCHI cells were cultured in RPMI 1640 supplemented with 10% bovine serum and 1× penicillin/streptomycin. All cells were maintained at 37℃ with 5% CO_2_ and passed using standard cell culture procedures.

### RNA interference and colony formation assay

FT246 (RRID:CVCL_UH61), FT282 (RRID:CVCL_A4AX), OVCAR4 (no RRID available) and KURAMOCHI (no RRID available) cells were reverse transfected with two independent non-targeting siRNAs (Horizon Discovery catalogue number: D-001810-10 and one custom control pool) or pooled MECOM, PAX8, SOX17 and WT1 oligonucleotides (Horizon Discovery catalogue numbers: L-006530-02, L-003778-00, L-013028-01, L-009101-00). Cells were seeded for colony formation assays as previously described (Reddy et al., 2021). Each assay was performed at least three times, taking the average of technical triplicate wells from each set.

### THZ1, THZ531 and JQ1 dose-response curves

FT246, FT282, OVCAR4 and KURAMOCHI cells were plated in 96 well plates and treated with THZ1 (Selleck Chemicals), THZ531 (ApexBio), JQ1 (Tocris) or dimethyl sulfoxide as a vehicle control. Each drug was added at 1:3 dilutions starting from 10,000 nM to 1.5nM as triplicates and incubated for 72 hours at 37℃. Cell survival was quantified with the Promega Cell Titer Glo reagent kit following manufacturers guidelines. Each assay was performed at least three times, taking the average of technical triplicates well from each set.

### RNA-seq differential and global expression analysis

FT246 and KURAMOCHI cells were reverse transfected with siRNAs for two scramble controls, *MECOM, PAX8, SOX17* or *WT1* as described above. Media were replaced 24 hours later. Total RNA was harvested from an equal number of cells 72 hours post reverse-transfection and spiked with an exogenous ERCC spike-in (Thermo Fisher). Non-stranded Poly-A libraries were constructed and sequenced with 150 base pair sequencing at 40 million reads on the DNB-seq DNA sequencing platform. Hisat2-2.1.0 was used for alignment with the reference genome gencodev26 + ERCC92. The count matrix of gene-level reads were generated using ht-seq count (v0.5.4.3). For DESEQ2 analysis, ERCC read counts were removed and gene expression analysis was performed with DESeq2 (version 1.24.0). For global ERCC spike-in analysis the transcripts per million were calculated from the count matrix followed by loess regression normalization using the *affy* package in R. Technical duplicate samples were generated on independent days.

### H3K27ac CUT&TAG in FTSECs and bioinformatics

CUT&TAG was conducted based on the CUTANA CUT&TAG protocol v1.5 (Epicypher). Isolated nuclei from primary fallopian tube secretory epithelial cells bound to concalavin A beads were incubated with primary antibody that detects H3K27ac (0.7μg, Diagenode, RRID:AB_2637079) at 4℃ overnight. On day 2 samples were incubated with secondary antibody and washed. Samples were then incubated with pAG-Tn5 for 1 hour, washed and placed in a tagmentation buffer followed by 1hr incubation at 37℃. Beads were then washed with TAPS buffer followed by incubation in SDS release buffer for 1 hour at 58℃. The reaction was quenched with SDS Quench buffer. PCR was conducted and DNA cleanup was performed with 1.3× Ampure beads. Samples were sequenced at the CSMC AGCT Genomics Core facility. Data processing was conducted following the workflow described by Zhen et al https://yezhengstat.github.io/CUTTag_tutorial/. Briefly, FASTQ files were aligned to hg38 with Bowtie. Duplicates and reads with alignment quality scores of 2 or lower were removed. Peaks were called using SEACR with the parameters set to --0.01 --norm -- stringent.

### CUT&RUN and bioinformatics

CUT&RUN was performed based on the CUTANA CUT&RUN protocol v1.6 (Epicypher). Isolated nuclei from FT282 and KURAMOCHI were bound to Concalavin A beads and incubated overnight at 4℃ with primary antibody for H3K27me3 (Cell Signaling, RRID:AB_2798370), H3K27ac (Diagenode, RRID:AB_2637079), MECOM (Cell Signaling, AB_2184098), PAX8 (Novus, RRID:AB_2283498), SOX17 (Abcam, RRID:AB_2801385), WT1 (Santa Cruz, RRID:AB_632611). Samples were washed after overnight incubation and treated with pAG-Mnase for 10 minutes at room temperature followed by activation by calcium chloride for 2 hours at 4℃. The reaction was quenched with STOP buffer and DNA was extracted with a DNA purification kit (Epicypher). Library preparation was conducted with the NEBnext Ultra II kit with the following protocol adjustments. The second incubation during the end prep was extended to 1 hour at 50 degrees and all SPRI select steps were conducted at 1.5× volume. Cycling parameters were set to parameters as instructed in the CUT&RUN protocol v1.6 (Epicypher). Processing and peak calls were made using the same pipeline as the CUT&TAG pipeline.

### Hi-C and bioinformatics

FT246, FT282, KURAMOCHI and OVCAR4 Hi-C were conducted and processed by Active Motif using the Arima-Hi-C kit and library prep performed with technical duplicates (Arima Genomics). Prepped libraries were sequenced to a depth of 1 billion reads. FASTQ files were processed with the Juicer software package with default parameters for the generation of intermediate .hic files. Loop and TAD calls were generated using HiCCUPs as described previously (Rao et al., 2014).

### Data availability

All data have been deposited into the Gene Expression Omnibus (https://www.ncbi.nlm.nih.gov/geo/). RNA-seq following transcription factor knockdown and siRNA control treatments are available accessions GSE151316 (FT282 and OVCAR4) and GSE227276 (FT246 and KURAMOCHI); CUT&RUN and ATAC-seq data are available under accession GSE227779.

### Materials availability

This study did not generate any new materials.

## ACKNOWLEDGEMENTS

This project was supported by an Ovarian Cancer Research Fund Alliance Liz Tilberis Early Career Award (599175) (to K.L.), an American Cancer Society Research Scholars Grant (134005-RSG-19-135-01-DMC to K.L.), and an Ovarian Cancer Research Fund Alliance Program Project Development award (373356) (K.L.). The research described was supported in part by NIH/National Center for Advancing Translational Science (NCATS) UCLA CTSI grant number UL1TR001881. R.N. is supported in part by a Ruth L. Kirschstein Institutional National Research Service Award (T32) from the NIH (grant number 5 T32 GM 118288-2) and UCLA CTSI core voucher number V175. The content is solely the responsibility of the authors and does not necessarily represent the official views of the NIH. This work used equipment and services provided at the Cedars-Sinai Biobank and Translational Research Core and the Applied Genomics, Computation and Translation (AGCT) Core. Hi-C data was generated and processed by services provided by Active Motif. The results shown here are in whole or part based upon data generated by the TCGA Research Network: www.cancer.gov/tcga and The Genotype-Tissue Expression: https://gtexportal.org/home/consortium.

